# Modulation of liposome membranes by the C-terminal domain of the coronavirus envelope protein

**DOI:** 10.64898/2026.03.23.713574

**Authors:** Reema Alag, Bui My Hanh, Ali Miserez, Jaume Torres, Konstantin Pervushin, Bhargy Sharma

**Affiliations:** School of Biological Sciences, Nanyang Technological University, Singapore; School of Materials Science and Engineering, Nanyang Technological University, Singapore

## Abstract

The coronavirus envelope (E) protein is a viroporin that plays a key role in viral assembly, release, budding and pathogenesis. E protein forms oligomeric ion channels that can activate immune responses. However, high-resolution structural data for its extramembrane domains is limited. The C-terminal domain of SARS-CoV has been shown previously to form amyloid fibers, and here we show that these fibers can modulate the shape of liposomes. The propensity to form fibrils, and their effect on liposomes, was examined for sequences belonging to the four clades of coronaviruses. Electron microscopy data shows that the C-terminal domain in E protein adopts a filamentous structure. These findings demonstrate the potential of these peptides to modulate membranes, providing a possible mechanism by which E protein interacts with membranes in the host cell.

## INTRODUCTION

Coronaviruses (CoVs) are a large family of viruses in the order *Nidovirales* and the family *Coronaviridae*. CoVs are classified into four genera: Alphacoronavirus (αCoV), Betacoronavirus (βCoV), Gammacoronavirus (γCoV) and Deltacoronavirus (δCoV), within the subfamily *Orthocoronavirinae*^1^. With over 40 CoV species categorised taxonomically, αCoVs and βCoVs circulate among mammalian hosts, while γCoVs and δCoVs primarily infect birds. CoVs are enveloped viruses characterised by a large, positive-sense, single-stranded RNA genome and four structural proteins: spike (S), matrix (M) protein, nucleocapsid (N) and envelope (E). These proteins play essential roles in virion assembly and infectivity.

The virus responsible for the COVID-19 pandemic, the Severe Acute Respiratory Syndrome coronavirus (SARS-2), belongs to the βCoV clade. Human CoVs (hCoVs) can cause respiratory and enteric diseases of varying severity. There are seven known hCoVs: 229E, NL63, OC43, HKU1, SARS and SARS-CoV-2, and Middle East respiratory syndrome coronavirus (MERS)^2^. Next generation sequencing has shown that SARS-2 has 79% homology to SARS and 50% homology to MERS^3^. The pandemic triggered fundamental studies into the structures and functions of the CoVs and their proteins^4^. Nearly a decade later, we possess detailed structural models of several key virus proteins^5–9^.

The envelope (E) protein is encoded by SARS coronavirus 2 (SARS-CoV-2), the agent that caused the recent COVID-19 pandemic, but also in many other pathogenic coronaviruses that infect humans and livestock, with prime economic importance. Therapeutic treatments for coronavirus infections are ineffective, thus there is an urgent global need for antivirals^10–14^ and novel targets. In infected cells, the E protein is found in the endoplasmic reticulum Golgi intermediate compartment (ERGIC)^15^ where it forms pentameric ion channels^16–24^ and participates in CoV assembly and budding. Therefore, it is a confirmed drug target^17,25–29^. E protein is one of the structural proteins that shows highly conserved sequences in all coronaviruses and consists of the N-terminal domain, transmembrane domain (TMD) and C-terminal domain ^29,30^. Virus lacking E protein shows a significant reduction in viral titer, which confirms the role of E protein in virus production and maturation^31^. Studies of SARS-CoV E protein have shown a higher level of E-protein expression in infected cells during the viral replication cycle^32,33^. The TMD of E-protein has been studied extensively as a viroporin with ion channel activity, through self-assembly into a pentameric structure^34^. High-resolution models of the pentamer formed by the single α-helical TMD are available^16,35–37^, solved by solid-state NMR (ssNMR). However, the function and structure of the C-terminal domain remain poorly studied. It is also not clear how the ion channel activity of the E-protein is responsible for its function of assembly and budding^38^. A transition from α-helix to β-structure is induced by oligomerization that is favored when protein concentration is higher (low lipid-to-protein ratios, LPR)^24^. ssNMR has shown that truncated E protein (residues 8-65) forms β-strands at LPR20^39^. The C-terminal domain is α-helical in isolated monomers^23,40^, but forms triple-stranded sheets which are interlocked outside the membrane^24,39^, likely triggered by oligomerization into pentameric or larger oligomers. This confirms previous experimental and in silico observations of a high tendency to aggregation and formation of amyloid fibers in this part of the protein^23,41^. This dramatic conformational change is intriguing since this predicted ‘beta-Proline-beta’ motif is found in all E proteins and thus it should have a functional role^42^. Oligomeric proteins have been previously reported to stabilize and modulate deformed liposome structures in the membranes^43–45^. Membrane curvature modulation through these protein assemblies is a common feature between virulent bacterial and viral infections. CoVs, like flaviviruses, noroviruses, and picornaviruses, induce the formation of double-membrane vesicles during RNA replication. These vesicles serve as scaffolds for viral replication, while other topological models in alphaviruses and nodaviruses induce membrane spherules for replication^46^. Biological membranes can adopt various shapes, with several morphologies observed in intracellular organelles. Local curvature at double membrane vesicle junctions influences the organization and function of associated pore complexes^47^. Detailed visualization of these assemblies has been limited due to low resolution hindering the identification of missing components and electron densities within these complexes. Nonetheless, symmetrical filamentous architectures are recurring features in CoV virion formation and replication^46^. Membrane modulation by amyloid fibrils has shown membrane curvature and rupture in the presence of amyloid fibrils^48^. Previous reports have identified self-assembling helical filaments in both bacterial and viral systems that exhibit potential gating mechanisms for protein translocation across membranes^49–52^. Structural organization is sensitive to amino acid composition; for example, β-sheet conformations promote lamellar morphologies, while α-helical segments stabilize rod-like assemblies^53^.

Here, we extracted E protein sequences from α, β, γ, and δ clades of coronaviruses and studied the formation of fibrils by the C-terminal residues of several CoV-E proteins from these four clades of coronaviruses. These fibrils demonstrate ability to deform the membrane bilayer of the intracellular ERGIC liposomes leading to variations in the local curvatures and their ability to deform the membrane is consistent throughout all the clades The interactions between the fibrils formed by C-terminal residues of CoV-E proteins and ERGIC liposomes are two-fold: the liposome increase the fibrillation, and the presence of fibrils, in turn, modulate the liposome shape and disruption. The C-terminal of the E protein has a Golgi-complex targeting signal and brings the E protein to the Golgi-complex^15^. We hypothesize that the amyloidogenic C-terminal region of the E protein fibrils modulates or ruptures the Golgi membrane. Our high-resolution cryo-EM data and biophysical assays demonstrate the formation of filamentous structures by E protein C-terminal residues, which consist of β sheets^39^.

## RESULTS AND DISCUSSION

### Amyloid-forming sequences are present in the C-terminus of CoV E proteins

Amyloid propensity is observed in the C-terminal residues of E-proteins across all clades^54^. Many of these sequences consist of aromatic and positively charged residues, which play an important role in amyloidogenic propensity^55,56^. From the Multiple Sequence Alignment (MSA) derived from 568 E protein sequences (218 from αCoVs, 126 from βCoVs, 195 from γCoVs, and 29 from δCoVs), representative peptides were selected for predicting amyloidogenic regions present in most species within each clade using Waltz^57^. Selected peptide sequences from βCoVs SARS (YP_009825054.1) and SARS-2 (YP_009724392.1), Mink αCoV (QKX95791.1), Porcine δCoV (AWV96567.1), and Avian γCoV (QYL35049.1) were aligned based on sequence homology (Figure 1). A total of six peptides were used in this study for fibrillation experiments, with one from each clade, and two different polypeptide lengths chosen for the γCoV sequence. (Supplementary Table 1).

**Figure 1.**
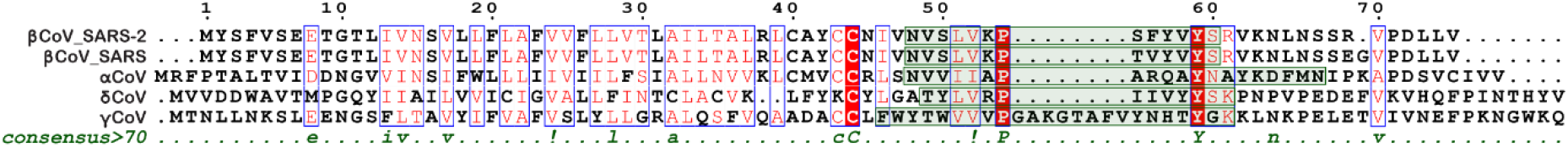
Sequence alignment of representative coronavirus E-protein sequences, including βCoV_SARS-2 (YP_009724392.1), βCoV_SARS (YP_009825054.1), Mink αCoV (QKX95791.1), Porcine δCoV (AWV96567.1), and Avian γCoV (QYL35049.1). Predicted amyloid-forming peptide regions are highlighted in green, and details are mentioned in Supplementary Table 1. Sequences were aligned using ClustalW. The consensus sequence and annotation were generated using ESPript. In the ESPript scheme, red boxes with white characters indicate strict identity, red characters denote similarity within a residue group, and blue frames indicate similarity across different residue groups.

### C-terminal tail sequences of E proteins form polymorphic fibrillar structures

First, we studied the fibrillation of the six peptides at pH 5.8, as shown in the Supplementary Figure 2. The peptide from αCoV formed networks of long, unbranched structures, while SARS-CoV-2 readily formed twisted fibril networks (Supplementary Figure 2A-B). The peptide from δCoV formed thin, long fibrils (Supplementary Figure 2C). The peptide from γCoV_14 formed straight, rigid fibrils (Supplementary Figure 2D). The longer peptide from γCoV_24 formed flexible, elongated structures rather than well-defined fibrils (Supplementary Figure 2E). The slightly acidic pH which mimics physiological acidic environments of organelles like endosomes, trans Golgi-network and inflamed tissues^58,59^. The kinetics of fibril formation were studied using the Thioflavin T (ThT) fluorescence assay. All peptides showed a concentration-dependent increase in the number of fibrils, but the minimal concentration required was clade-dependent (Figure 2). For αCoV, SARS-CoV-2, and γCov_24, the minimum concentration needed for the ThT assay was 0.1 mM, while δCoV showed very low fibrillation at 0.1 mM after a long incubation. For αCoV peptide, there was a lag of over 40 h at 0.1 mM, but this lag time was reduced to half at 0.2 mM (Figure 2A). The SARS peptide readily showed fibrillation at 0.1 mM and 0.2 mM, with higher initial fluorescence at 0.2 mM and a slight lag time (Figure 2B). SARS-2 readily formed fibrils with no lag at both 0.1 mM and 0.2 mM (Figure 2C). The peptide from δCoV exhibited slow fibrillation at 0.1 mM, starting after 60 h of incubation, whereas rapid fibrillation was observed at 0.3 mM, reaching saturation within 30 h (Figure 2D). This sample at 0.3 mM was later used for cryoEM structure determination. However, γCov_14 did not exhibit fluorescence at 0.1 mM; the minimum concentration required for the ThT assay was 0.4 mM. The peptide from γCoV_14 formed straight and rigid fibrils (Figure 2E) and required 0.4 mM to show fibrillation kinetics. A sigmoidal growth curve was observed at 0.5 mM. The longer peptide from γCoV_24 showed rapid fibrillation obtained at 0.1 mM and 0.2 mM (Figure 2F). Various factors influence the initial peptide concentrations required for amyloid formation, including the Critical Aggregation Concentration, an inherent property of the peptide^60^. Lower concentrations of the peptides are insufficient to produce enough fibrils to reach the detection limit of ThT, affecting the aggregation kinetics^61^. The binding affinity and stoichiometry vary, causing some peptides to have greater affinity for ThT or more binding sites than others.

**Figure 2.**
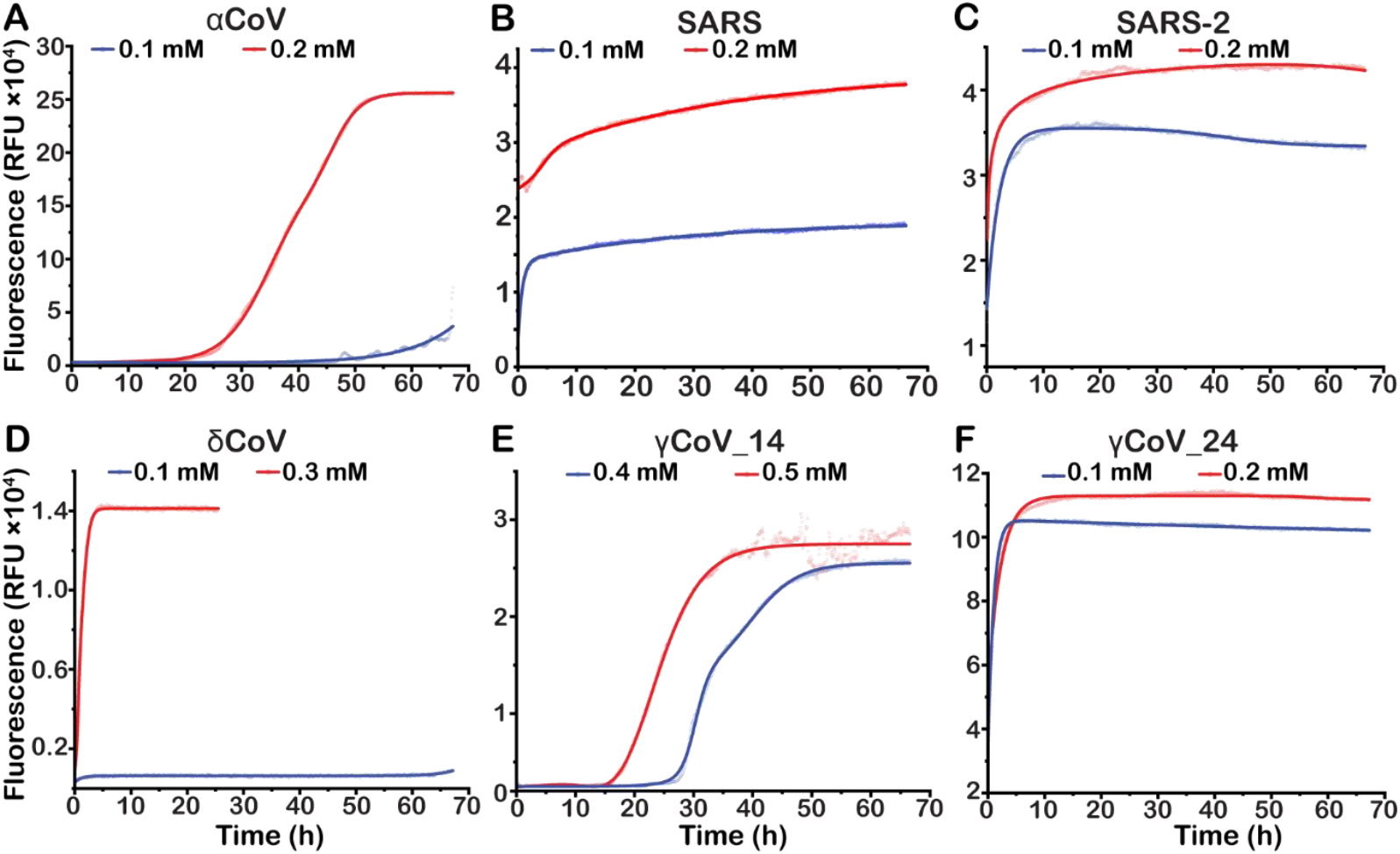
Thioflavin T (ThT) fluorescence assays showing fibrillation kinetics of (A) αCoV, (B) SARS, (C) SARS-2, (D) δCoV, (E) γCoV_14 and (F) γCoV_24 peptides at lower and higher concentrations are represented by blue and red curves, respectively. ThT fluorescence intensity increases with peptide concentration, consistent with enhanced fibril formation. The experimental data points were fitted using dose-response curves represented as solid lines using OriginLab software. Fluorescence values for buffer controls have been subtracted from each curve.

At physiological pH 7.5 (Figure 3), αCoV showed faster kinetics, with a lag phase of 16 hat 0.1 mM and 4 hours at 0.2 mM (Figure 3A). SARS formed fibrils at both 0.1 mM and 0.2 mM (Figure 3B), and SARS-2 fibrils displayed rapid kinetics at these concentrations (Figure 3C). At pH 7.5, δCoV did not exhibit fibrillation at 0.1 mM, but showed faster kinetics at 0.3 mM peptide concentration compared to pH 5.8 (Figure 3D). At pH 7.5, γCov_14 had a longer initial lag phase but faster kinetics at both 0.4 mM and 0.5 mM (Figure 3E). γCov_24 displayed faster kinetics at pH 7.5 compared to pH 5.8 for 0.1 mM and 0.2 mM peptide (Figure 3F). We posit that the pH changes influence peptide protonation, which in turn affects attraction and repulsion between peptides, thereby impacting aggregation and reaction rates^62^. At physiological pH 7.5, differences in fibril morphology were also observed (Supplementary figure 3). Fibrils from αCoV E were straighter than those formed at acidic pH (Supplementary Figure 3A). SARS fibrils appeared wider and straighter (Supplementary Figure 3B), while SARS-2 formed polymorphic fibrils (Supplementary Figure 3C). γCov_14 formed straight, branched fibrils, but no fibrils were seen in the longer γCov_24 sequence (Supplementary Figure 3D,E). At both acidic and physiological pH, TEM micrographs and ThT assays reveal an innate ability of these E peptides to form amyloid-like fibrils under physiological and subcellular conditions, albeit with different structures and kinetics, suggesting distinct nucleation mechanisms^63^.

**Figure 3.**
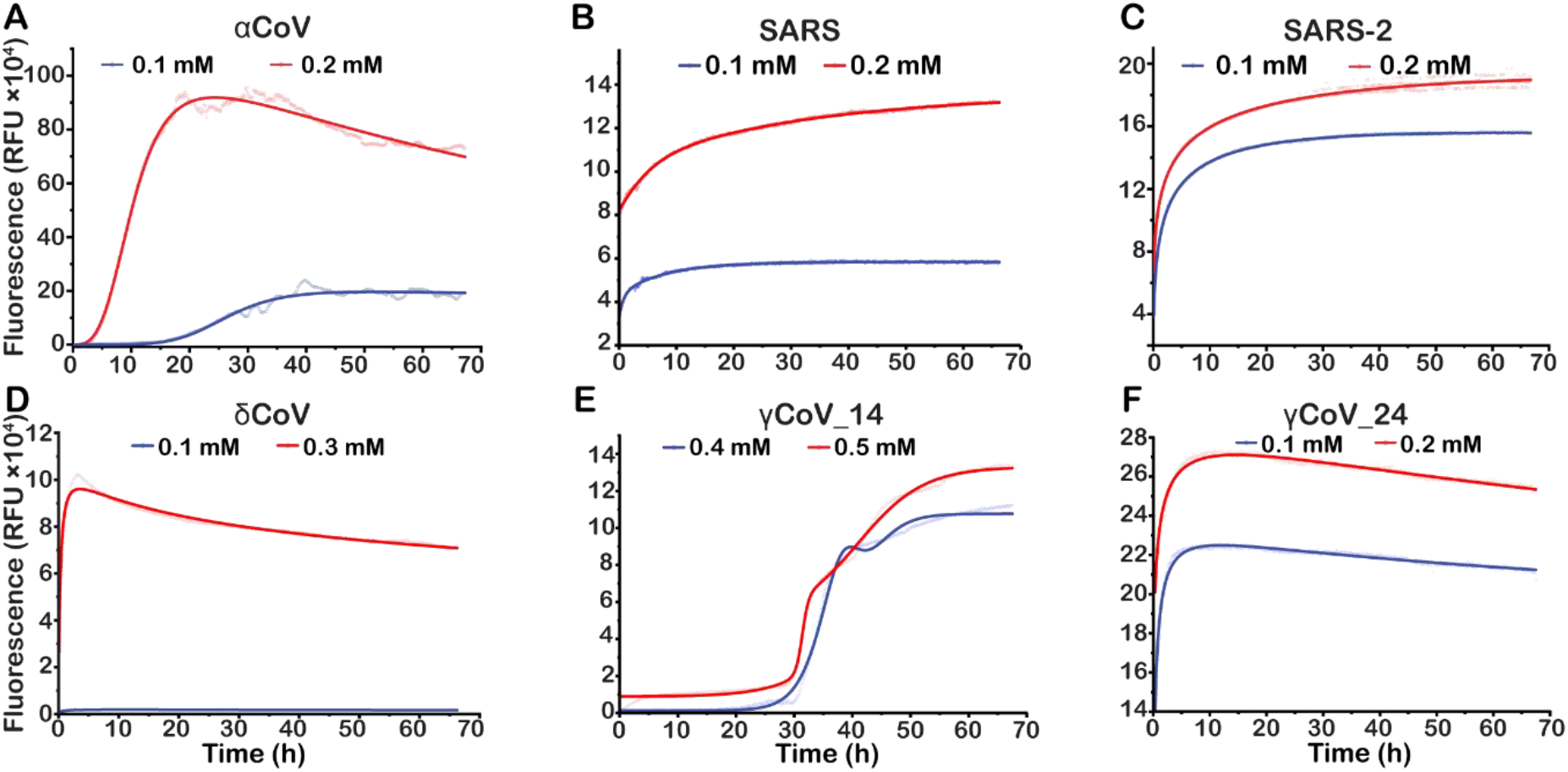
Thioflavin T (ThT) fluorescence kinetics for fibrillation of (A) αCoV, (B) SARS, (C) SARS-2, (D) δCoV, (E) γCoV_14 and (F) γCoV_24 peptides at lower and higher concentrations are represented by blue and red curves, respectively. ThT fluorescence intensity increases with peptide concentration, indicating enhanced fibril formation. The experimental data points were fitted using dose-response curves represented as solid lines. Fluorescence values for buffer controls have been subtracted from each curve.

### CoV E peptide fibrils can modulate ERGIC-like membranes

To study the effect of amyloids on ERGIC-like membranes as well as to see the ergic-liposome and amyloid interaction effect in all four clades, we chose one peptide from each clade, including αCoV, βCoV_SARS2, δCoV, and γCoV_24. Fibrils for all four peptides were grown in the presence of liposomes at Liposome-to-Peptide ratios (herein, L:P) 0.5:1 and 1:1 at 37º and 180 rpm in incubator shaker at pH5.8. Only liposome was used as control and incubated at same conditions as fibrils. The interaction between E-derived fibrils and ERGIC liposomes were imaged through wide-field negative-stained TEM micrographs (Figure 4). Only liposomes, in the absence of fibrils, show circular morphology as expected (Figure 4A). However, in the presence of fibrils at L:P of 0.5:1 it was observed that the presence of fibrils changed the liposome shape from smooth circular to sharp-edged polygonal structures (Figure 4B). It has been found that polyhedral shape of liposomes are associated with a higher degree of disorder and curvature and difference in liposome morphology might be associated with membrane bending forces^64^. It has been reported that polyhedral shape of liposomes helps in endosomal escape mechanisms which involved pore formation, rupture and membrane fusion^65^. It is well documented that curved structures like polygonal structures, cubic phases and other disordered structure of membranes are found in living cellular systems. Sometimes these structures are found on a small and local scale specially in context of fusion and fission processes^66^. Previous studies with Islet amyloid polypeptide (IAPP) and phospholipid vesicle interaction have shown that IAPP induces curvature and disruption in vesicle membrane^48^. At L:P ratio of 1:1, TEM images of αCoV, δCoV and γCoV_24 peptides with liposomes showed the mixture of polygonal liposome and ruptured liposome, indicating that higher L:P ratio can rupture the lysosomes. However, for βCoV_SARS2 at L:P of 1:1, polygonal liposomes were not present and only some ruptured liposomes were seen (Figure 4C and Supplementary Figure 4). It has been reported previously for Aβ and β2-microglobulin fibrils can ruptured the liposomes membranes^67,68^. Our results demonstrate that as C-terminal of E-protein make fibrils and these fibrils can modulate the membrane therefore the C-terminal peptide can modulate or bend the membrane and might responsible for assemble and budding function of E-protein. Our results are consistent with the recent finding about function of C-terminal of E-protein which shas shown that amyloid making motifs of Sars-Cov2 is responsible for VLP (virus like particles) release and mutation in F56/Y57/Y59 residues are critical for E-protein function in VLP release^38^.

**Figure 4.**
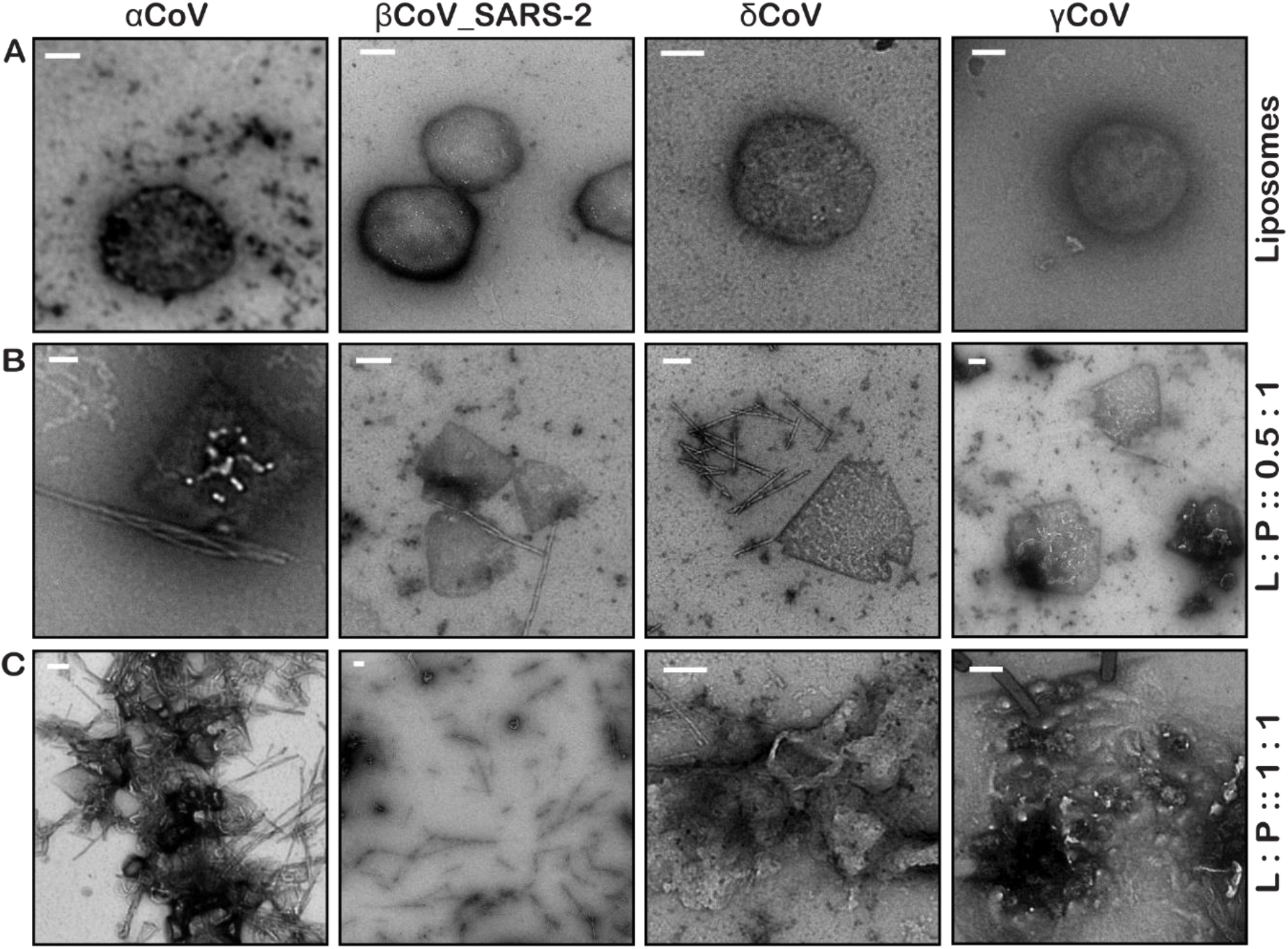
Transmission electron micrographs showing effect of CoV peptide fibrils on morphology of liposomes. (A) TEM images of liposomes with circular morphology in control sample versus polygonal shape in presence of αCoV peptide in 0.5:1 ratio. (B) TEM images of nearly circular liposomes changing to triangular morphology in the presence of peptide fibrils at L:P ratio 0.5:1. (C) TEM images of disrupted liposomes at an L:P ratio of 1:1. Scale bars in all images represent 200 nm.

### Liposomes impact the fibrillation of the C-terminal peptides

To investigate the impact of liposomes on the fibrillation behaviour of the CoV peptides, we studied fibril growth kinetics at L:P ratios 0.5:1 and 1:1 using the ThT assay (Figure 5). For the C-terminal peptide derived from αCoV E, the presence of liposomes halved the lag phase duration, from 20 h for the peptide alone, to 10 h in the presence of liposomes at L:P 1:1 (Figure 5A). For SARS-2 peptide, at an L:P ratio of 0.5:1, fibril growth was not affected by liposomes, but at an L:P ratio of 1:1, it increased the number of fibrils (Figure 5B). For the δCoV E peptide, fibril formation was lipid-dependent (Figure 5C). For the γCoV-24 E peptide, the fluorescence was unchanged at L:P ratios 0.5:1 and 1:1 (Figure 5D). Our results indicate the increase in fibril formation in the presence of liposomes which is consistent with previous studies showing an increase in fibril formation in the presence of liposomes^69^.

**Figure 5.**
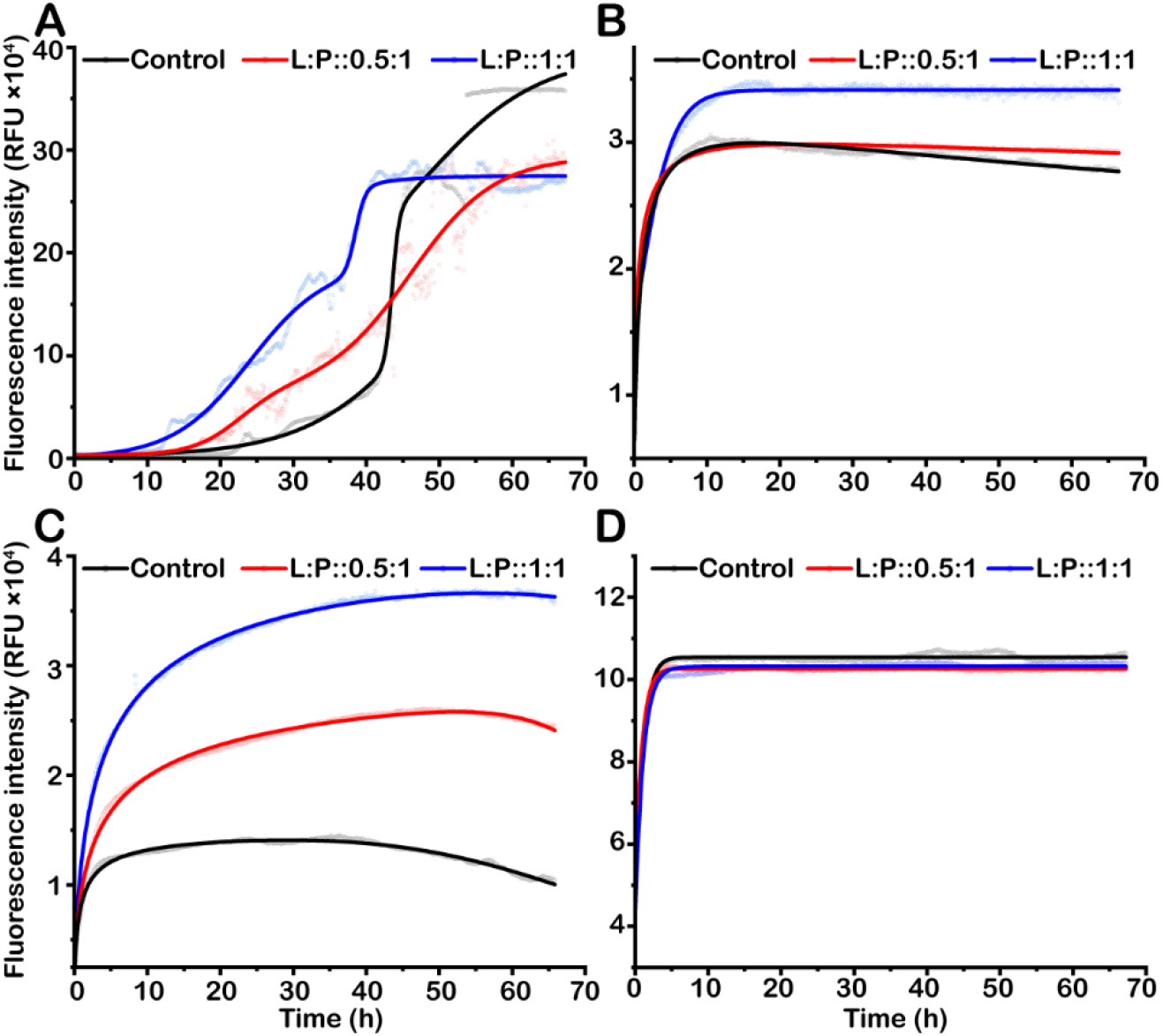
Thioflavin T (ThT) fluorescence assay for (A) αCoV, (B) βCoV-2, (C) δCoV, (D) γCoV-24, exhibiting kinetics of fibril formation. The fibrils growing in the absence of liposomes are shown as control (black), with L:P 0.5:1 and 1:1 are represented shown in red and blue, respectively. The experimental raw data points were fitted using dose response curves represented as solid lines. Fluorescence values for buffer controls have been subtracted from each curve.

**Figure 6.**
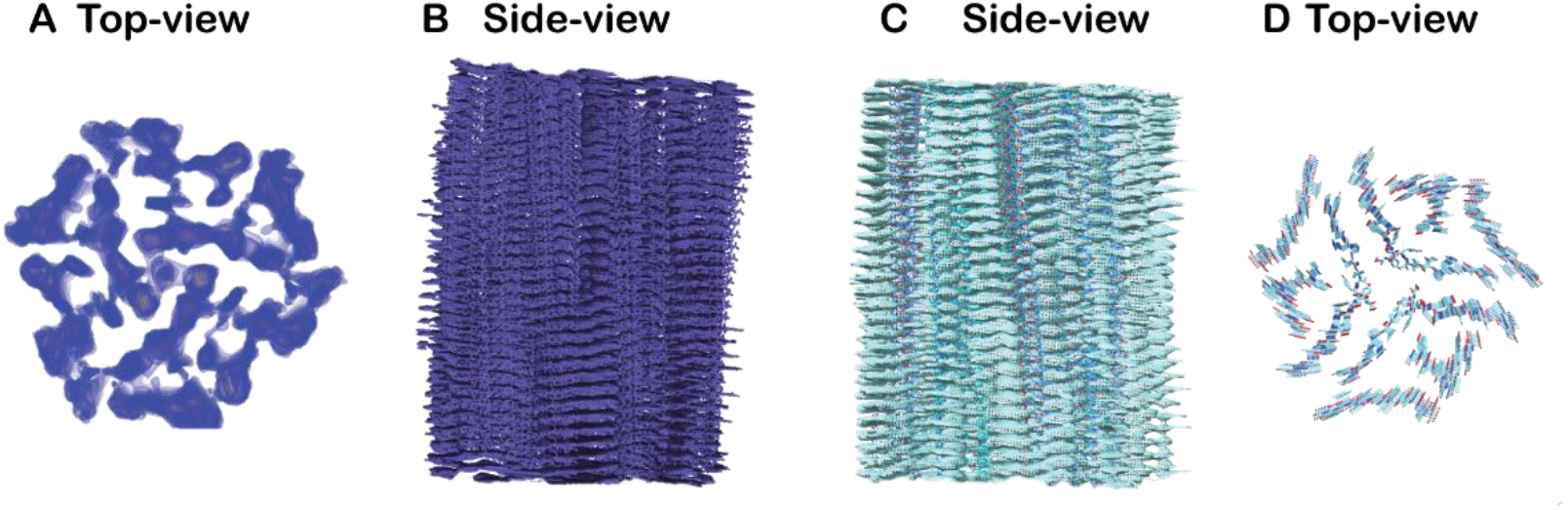
3D reconstructed map and atomic model of the δCoV fibril. (A) Top view and corresponding (B) side view of the reconstructed δCoV fibril density map. (C) Side view of the refined atomic model generated using Phenix, Coot and Chimera, (B) Corresponding top view of the atomic model.

### CoV-E peptide fibrils are comprised of mixed secondary structures

For the prediction of the fibril structure, 10 units of CoV-E peptides were input in each query in AlphaFold^70^. AlphaFold predicted a β-sheet fibril model for SARS, SARS-2, δCoV, and γCoV-24 peptides, a very low confidence fibril model for γCoV-14, and a multimeric model with a pore for the αCoV (Supplementary Figure 5). The fibrils showed structural polymorphism at different concentrations, as apparent from the negative stained TEM images. We used circular dichroism (CD) to determine the secondary structures in mature fibrils at 0.1 mM after a one-week incubation (Supplementary Figure 6). The secondary structure was obtained using BestSel^71^ and K2D2^72^ algorithms (Supplementary Tables 2 and 3). However, the maximum error for predicted composition through K2D2 exceeded 0.4 for the fibrils grown at pH 7.5. Additionally, due to the polymorphic nature of these fibrils, the contributions of each secondary structure varied slightly between different batches. We used Fourier transform infrared spectroscopy (FT-IR) to evaluate the secondary-structure composition of fibrils grown from 0.2 mM peptides at both pH values^73,74^. The amide I peak was deconvoluted to identify the secondary-structure contributions of each component (Supplementary Figure 7). While the exact predicted contributions of the different secondary structures vary between the results of CD and FT-IR, both results were consistent with a mix of secondary structures within the peptide samples with an increasing contribution of ordered structures at pH 7.5 compared to pH 5.8 (Supplementary Table 4).

### Three-dimensional structure of δ-CoV fibrils shows cross-β structure

We studied the δCoV fibril structure at high resolution using cryo-electron microscopy (cryoEM), due to its smaller peptide length along with the presence of aromatic and charged residues. For cryoEM, 0.3 mM δCoV peptide was incubated for a week in fibrillation buffer at pH 5.8. CryoSPARC was used for data acquisition, processing and 3D map reconstruction (Supplementary Figure 8). Micrographs with individual fibrils with least overlapping regions were used for picking particles and 2D classes with no overlaps were selected for model building. The refinement was done in multiple iterations to get the best resolution model from the cryoEM. The best refinement revealed the formation of multi-stranded helical filaments at 3.6 Å, with a helical rise of 4 Å around the central axis in cross-β stacking (Supplementary Figure 9). The fibril showed a C3 symmetric architecture with a twist of 119.3°, indicating three subunits per helical turn. We identified four asymmetric units in the repeating lattice represented by the helical symmetry order of four. The model building from the 3D map and validation using Phenix confirmed the symmetry of the structure. The overall structure of the fibrils showed helices in the core interspersed with β structures and unstructured loops (Figure 7).

The presence of other secondary conformations in the fibril solution, as suggested by CD and FT-IR, indicates the dynamic process of fibrillation and kinetics in peptide solutions. However, through the selection of homogeneous structures within the electron micrographs of δCoV fibrils, we were able to show that the mature fibrils acquire the typical cross-β structure of amyloids.

## CONCLUSION

Our results provide mechanistic insights into fibrillar amyloids formed by the C-terminal residues of E proteins. Our findings reveal the crucial role of these polypeptide sequences in affecting membrane curvature and disrupting membranes in a concentration-dependent manner. We demonstrate the potential of peptides as membrane-modulating biomaterials. These C-terminal regions are promising targets for antiviral strategies, serving as key mediators of molecular interactions between the virus and host cells.

## MATERIALS AND METHODS

### Extracting the sequences

A basic local alignment search tool (BLAST) for the E protein from the four CoV genera was conducted via NCBI^75^. Approximately 5000 sequences of CoV genomes from the four genera were retrieved from NCBI. Genetic filtering and clustering were carried out to remove low-quality sequences to shortlist a good representation of sequences for each genus. As a result, a total of only 567 sequences for E protein (218 αCoV, 126 βCoV, 195 γCoV and 28 δCoV) represented by their GenBank Accession numbers, were retrieved in FASTA format, and used for analysis.

### Multiple Sequence Alignment of derived sequences for all the clades and amyloid predication for the sequences

218 sequences from α clade, 126 sequences from β clade, 195 sequences from γ clade and 28 sequences from δ clade were aligned according to their clade. All the sequences derived for different clades were aligned using Clustal Omega alignment^76,77^. All the selected sequences from sequence alignment results were screened for their amyloid making possibilities using predictions software Waltz^57^.

### Peptide design and synthesis

Based upon the multiple sequence alignment of each clade, region after the TMD was carefully studied and suitable sequences with higher probability of amyloid making were selected from each clade. We have selected SARS sequence (45-61): NIVNVSLVKPTVYVYSR; SARS-2 sequence (45-61): NIVNVSLVKPSFYVYSR from β clade, from α clade: accession number MN535737(51-71): NVVIIAPARQAYNAYKDFMN, from δ clade: accession number MF642323 (49-61): TYLVRPIIVYYS, from γ clade: accession number MZ325299 (47-70):LFWYTWVVVPGAKGTAFVYNHTYG and MZ325299 (57-70): GAKGTAFVYNHTYG sequences were selected for further studies. Above mentioned synthetic peptides were purchased from GenScript Biotech (Singapore) Pte Ltd. Along with Waltz prediction, precautions were taken while designing peptide in terms of net charge and presence of excess hydrophobic residues as they might hinder the synthesis of peptides.

### Preparation of monomeric peptides and setting up amyloid fibrils

To set up amyloids, it is necessary to start the process with monomeric peptides. In order to get monomeric peptides, lyophilized peptide powder was dissolved in HFIP(1,1,1,3,3,3-hexafluoro-2-propanol) with 1-1.5 mg/ml concentration depending upon the solubility of peptide and incubated for 20-30 minutes at room temperature until it becomes clear solution. HFIP dissolved peptide was aliquoted in eppedorf tubes, where each tube contains 0.5 mg of dissolved peptide. These tubes were freeze dried and dried peptide film containing 0.5 mg of peptide was obtained. Dried film was dissolved in DMSO with final concentration of 3.5 mM and prepared stock solution. The stock solution was prepared fresh every time before starting amyloid formation. Before setting up the fibrils, stock solution was treated in water bath sonicator for 3 times with 10 seconds each time at strong pulse. To set up the fibrils, stock solution was diluted in 20 mM phosphate buffer and 140 mM NaCl pH (7.5) or 20 mM phosphate buffer and 140 mM NaCl pH (5.8) with final concentration of 0.1 mM and 0.2 mM. Amyloid fibrils were allowed to grow at 37 degree and 180 rpm.

### Transmission Electron microscopy analysis of amyloid and Negative staining of the samples

For preforming TEM analysis of amyloids, sample was prepared using negative staining where 4 ul of the amyloid sample were applied onto the glow-discharged 400-mesh-size carbon coated copper grids for 1 minute followed by staining with 2%(v/v) uranyl acetate for 1minute. Surplus sample and staining solution were removed by filter paper (Whatman). Amyloid analyses were performed with transmission electron microscopy (TEM) with a FEI T12 transmission electron microscope equipped with a 4K CCD camera.

### ThT assay to monitor fibril formation

ThT which is a fluorophore and specifically binds to the β-sheet structures of amyloids and enhanced the fluorescence, was purchased from Sigma-Aldrich (Sigma-Aldrich Inc., MO). ThT was dissolved in 20 mM Phosphate buffer, 140 mM NaCl pH 7.5 or in 20 mM Phosphate buffer, 140 mM NaCl pH 5.8 depending upon the experiment. Final concentration of ThT was kept at 20 uM where as final conncetration of peptides were used from 0.05 mM to 0.5 mM depending upon the requirement of each peptide to make fibrils. To check the effect of liposome on fibril formation, fluorescence reading from liposome alone was subtracted from the liposome with fibril. In order to perform the assays peptides were dissolved in 20 mM phosphate buffer, 140 mM NaCl pH 7.5 or in 20 mM phosphate buffer, 140 mM NaCl pH 5.8 with different final concentration from 0.05 to 0.5 mM as mentioned earlier. Peptides were used from freshly prepared DMSO stock as mentioned above whereas final concentration of ThT was kept at 20 uM. To check the effect of liposome on amyloid formation, fibril formation was studied in the presence of liposome with liposome to peptide ratios of 0.05:1 and 1:1. ThT reaction mixture was prepared in a 96-well flat bottom polystyrol microplate (Greiner, Frickenhausen Germany) and plate was sealed to stop any evaporation. Measurements were taken in triplicates using a Cytation microplate reader where readings were taken at excitation wavelength at 440 nm and emission at 495 nm and orbital shaking were done before each measurement.

### Liposome preparation

ERGIC-mimetic membrane contained 1-palmitoyl-2-oleoyl-glycero-3-12 phosphocholine (POPC),1-palmitoyl-2-oleoyl-sn-glycero-3-phosphoethanolamine(POPE),bovine phosphatidylinositol (PI), 1-palmitoyl-2-oleoyl-sn-glycero-3-phospho-L-serine (POPS),and cholesterol (Chol). The POPC: POPE: PI: POPS: Chol were used with molar ratios of 45: 20: 13: 7:15. All lipids were purchased from Avanti Polar Lipids. To prepare liposomes stock with 7 mM concentration,1.5 mg of ERGIC lipids dissolved in chloroform was taken in glass vial. Chloroform was removed under gentle stream of nitrogen gas and lipid film was dried in vacuum chamber at room temperature overnight. The dried film was dissolved in 300 ul of 20 mM phosphate buffer, 140 mM NaCl (pH 5.8) lipid suspension was vortexes for 10 minutes to make it homogenous. Homogenous suspension was put in water bath sonicator for 1 minute at normal wave using probe sonicator followed by 5 minutes vortex, this step was repeated 3 times. The suspension was transferred in eppendorf tube and performed 7 rounds of freeze-thaw cycle at 42°C and liquid nitrogen.

### Cryo-EM grid preparation and data collection

4 μl of the 7 days grown δCoV fibrils (0.3 mM monomeric concentration before fibrillation) were dispensed on a fresh plasma-cleaned Quantifoil™ copper grid (1.2/1.3, 300 mesh). Cryo-grid preparation was done using Vitrobot Mark IV instrument (FEI/Thermo Scientific, USA) with single blot time 2.5 sec, blot force 5, wait time 5 sec, at 100 % humidity and 4 °C. Data were collected using a FEI Titan Krios 300 kV Cryo-TEM equipped with Falcon 4i direct electron detectors and a Selectris X energy filter. A total of 6160 movies were collected using the EPU software at a magnification of 108kx where the pixel spacing is 0.97 Å/pix. The exposure time for each movie was 1 sec with 40 frames and a total dose of 40 e-/Å2. Data were collected using a defocus range of -2 to 2 µm.

### Cryo-EM data processing and analysis

The movies were motion-corrected using patch motion correction in cryoSPARC live^78^. The CTF estimation was performed using Patch CTF estimation in cryoSPARC live. A total of 6051 micrographs passed curation and were used for further processing. Initially, 204 particles were manually picked from a subset of 30 micrographs. These particles were used for 2D classification, and the selected classes were further used as a reference for filament tracing for the complete curated dataset. After pick inspection, different extraction box sizes and masking diameters were used to obtain single fibrils without overlap. 2D classifications were performed iteratively, and good 2D classes containing approximately 219,882 particles were subjected to initial helical reconstruction without applying symmetry. Symmetry search was used to identify symmetry in the reconstruction, and helical refinement was performed based on the local-minima results. The local resolution map was estimated in cryoSPARC, with GSFSC resolution of 3.55 Å^79^. 3D classification without alignment was performed after homogeneous refinement to further prune the number of particles. The map was sharpened, and the model was generated using PHENIX, with the initial model used for model building in Coot^80^, based on the previously reported protocol for amyloid structure determination^81^. The final real-space refinement and validation were performed in the PHENIX suite^82,83^.

### Circular dichroism

CD spectrum for δCoV after 7 days of fibrillation over 200-250 nm was acquired using Chirascan (Applied PhotoPhysics). The sample was diluted 100 times in water to ensure sample absorbance was ensured to be <2 at all wavelengths to produce reliable values. Each collection was repeated over three scans, with a 2 sec averaging time at each wavelength, using a quartz cuvette with a 0.1 mm path length.

### Fourier transform – infrared spectroscopy

ATR-FTIR spectroscopy of 0.2 mM E-peptide samples was performed on a Bruker Vertex 70 (Massachusetts, USA) equipped with a PIKE Technologies MIRacle attenuated total reflection (ATR) ZnSe-Diamond 3-reflection accessory and an LN2-cooled MCT detector. Scans were obtained at ambient temperature over the range of 4000–750 cm−1 with a resolution of 2 cm−1, averaged over 128 scans. All spectra were processed in OPUS 6.5, in the sequence of water vapor subtraction, baseline correction, and normalization using the amide I band. The amide I band was deconvoluted by secondary derivation, with peak fitting performed using 100% Gaussian curves, keeping the individual FWHMs relatively consistent. The deconvoluted peaks were assigned to β-sheet, helix, and turns structures.

## CONTRIBUTIONS

R.A. conducted TEM and ThT experiments and prepared an initial draft and revised the manuscript. B.S. conducted cryoEM and CD experiments, prepared figures, drafted, revised, and edited the manuscript. B.M.H. conducted FTIR experiments. A.M. supervised the experiments. J.T. supervised the project and advised on experiments and reviewed the manuscript. K.P. got funding, supervised the study, advised on experiments, and reviewed the manuscript. All authors read and commented on the final version.

## ACKNOWLEDGEMENTS

The authors thank Dr. Wahyu Surya for guidance on liposome preparation and Dr. Wuan Geok Saw and Dr. Bich Ngoc Ann Tran from the NISB cryo-EM facility for their guidance on data acquisition and processing. We thank the Singapore Ministry of Education for funding under the grant MOE-T2EP30220, awarded to Profs Konstantin Pervushin and Jaume Torres as PI and co-PI, respectively.

## DATA AVAILABILITY STATEMENT

Data will be made available on request.

## Notes

### Competing Interest Statement

The authors have declared no competing interest.

