## Supplementary information for "Modulation of liposome membranes by the C-terminal domain of the coronavirus envelope protein"

### ***Liposome membrane modulation by C-terminal fibrils of E-protein in coronavirus***

*Supplementary Table 1. C-terminal sequences from CoV-E identified to form amyloids (sequences marked in green in Figure 1)*

| Sequence | aa | Organism | Clade | Accession No. |
|---|---|---|---|---|
| $^{51}$NVVIIAPARQAYNAYKDFMN$^{71}$ | 20 | Mink CoV | αCoV | Genbank: MN535737 (QKX95791.1) |
| $^{45}$NIVNVSLVKPTVYVYS$^{60}$ | 17 | SARS | βCoV | Genbank: NC_004718 (YP_009825054.1) |
| $^{45}$NIVNVSLVKPSFYVYS$^{60}$ | 17 | SARS-2 | βCoV | Genbank: NC_045512 (YP_009724392.1) |
| $^{49}$TYLVRPIIVYYS$^{61}$ | 12 | Porcine CoV | δCoV | Genbank: MF642323 (AWV96567.1) |
| $^{57}$GAKGTAFVYNHTYG$^{70}$ | 14 | Avian CoV | γCoV | Genbank: MZ325299 (QYL35049.1) |
| $^{47}$LFWYTWVVVPGAKGTAFVYNHTYG$^{70}$ | 24 | Avian CoV | γCoV | Genbank: MZ325299 (QYL35049.1) |

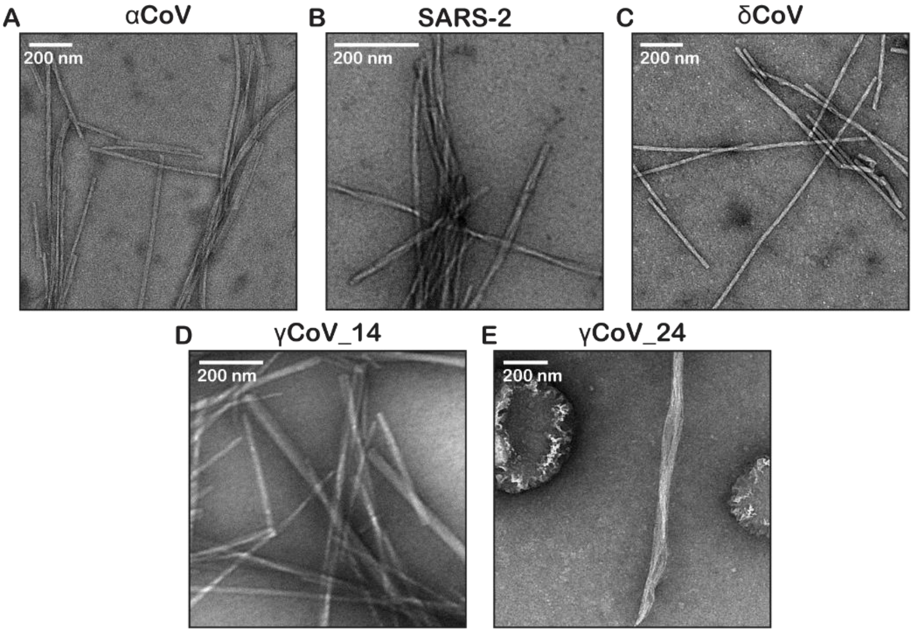


*Supplementary Figure 2. Structural and biophysical analysis of CoV peptides forming fibrils at pH 5.8. (A-E) Transmission Electron Microscopy (TEM) images of fibrils formed by (A) αCoV, (B) SARS-2, (C) δCoV, (D) γCoV_14 and (E) γCoV_24 peptides. All scale bars represent 200 nm.*

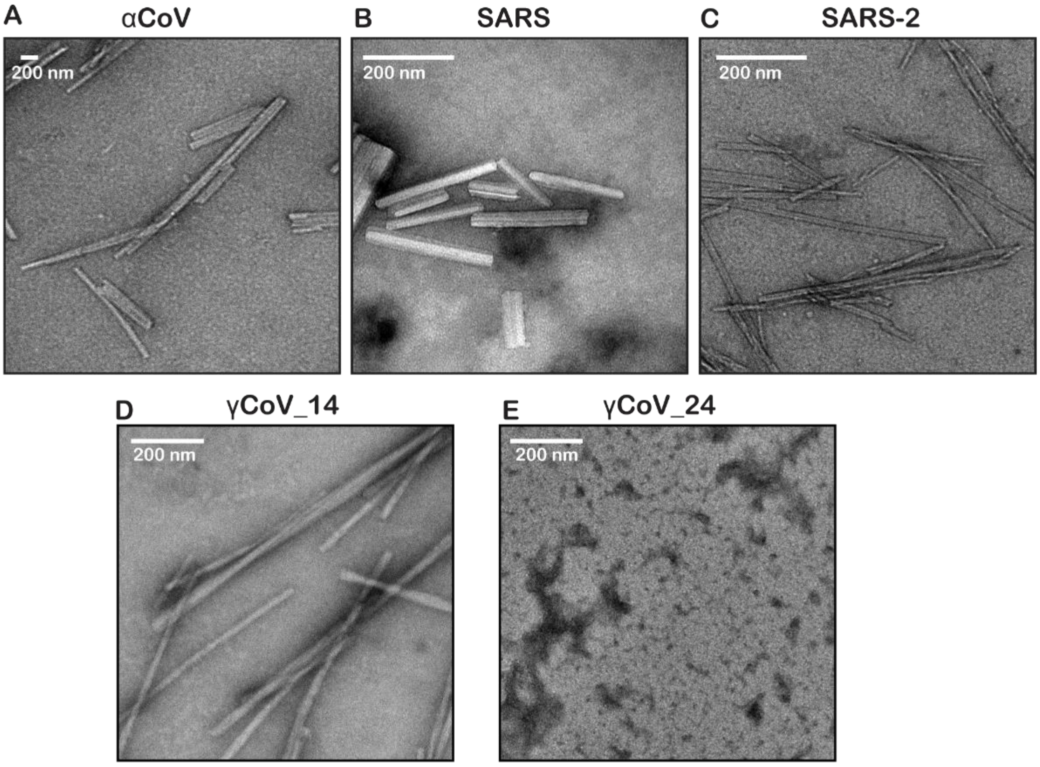


*Supplementary Figure 3. Fibril formation by CoV peptides at pH 7.5 indicated by TEM and ThT fluorescence assays. (A–C) Transmission electron microscopy (TEM) images of fibrils formed by E-protein peptides from (A) αCoV, (B) SARS, and (C) SARS-2. Scale bars: 200 nm.*

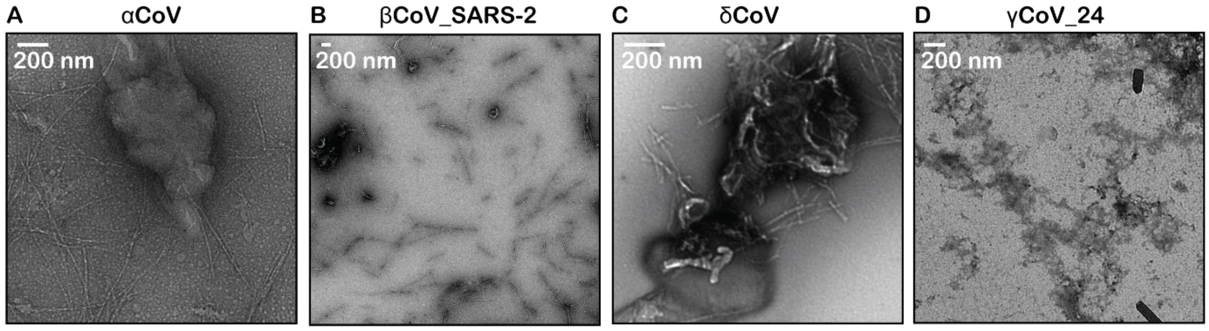


*Supplementary Figure 4. Higher fibril concentrations of (a) αCoV, (b) SARS-2, (c) δCoV, (d) γCoV-24 rupture the liposomes. Scale bars represent 200 nm.*

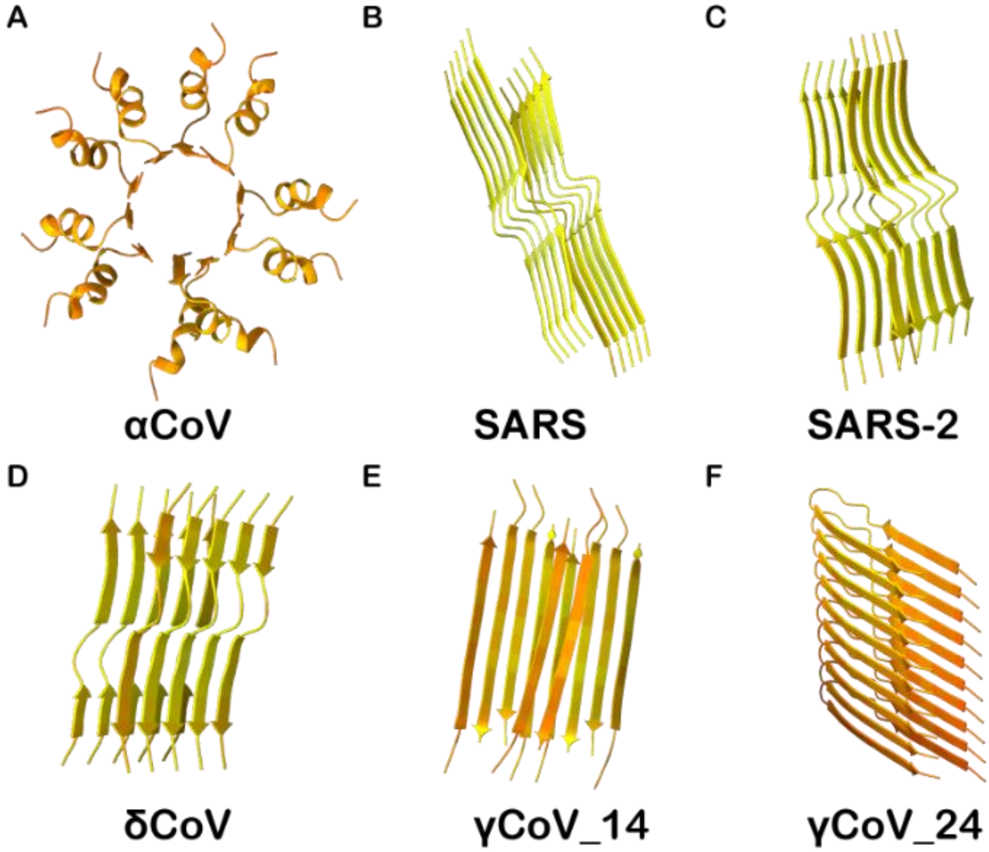


*Supplementary Figure 5. Alphafold structure prediction of multimeric C-terminal peptides. The colors represent confidence level as per AlphaFold pLDDT scores. (Orange represents pLDDT <50, yellow represents 50<pLDDT<70).*

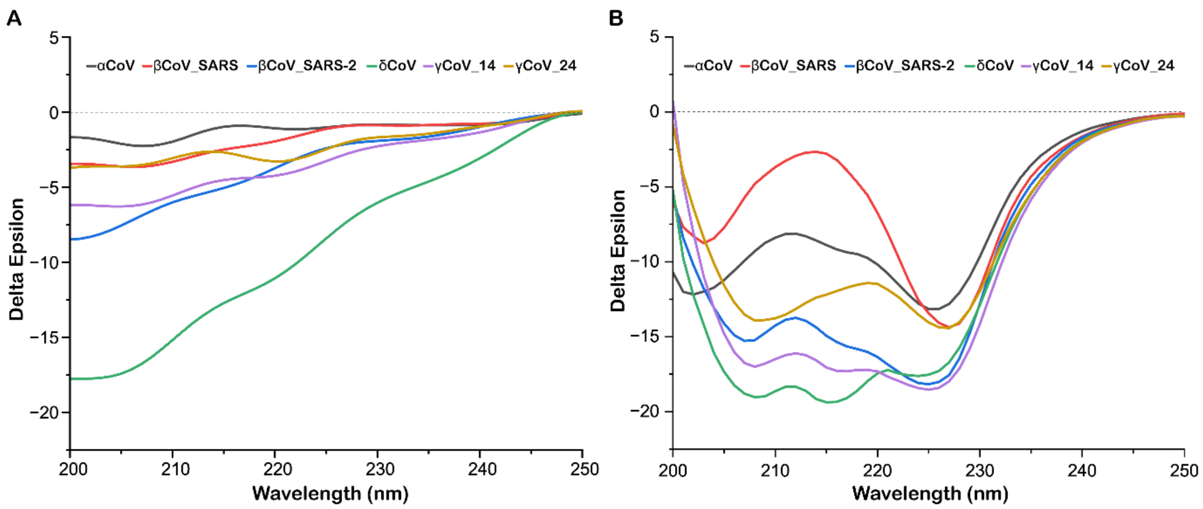


*Supplementary Figure 6: Circular dichroism spectra of the peptides grown for 1 week under fibrillation conditions at (a) pH 5.8 and (b) pH 7.5.*

*Supplementary Table 2: Secondary structures identified using CD in peptides grown for 1 week under fibrillation conditions at pH 5.8 and pH 7.5.*

| | αCoV | | βCoV-1 | | βCoV-2 | | δCoV | | γCoV_14 | | γCoV_24 | |
|---|---|---|---|---|---|---|---|---|---|---|---|---|
| | pH 5.8 | pH 7.5 | pH 5.8 | pH 7.5 | pH 5.8 | pH 7.5 | pH 5.8 | pH 7.5 | pH 5.8 | pH 7.5 | pH 5.8 | pH 7.5 |
| Helix | 43.7 | 55.9 | 28.6 | 14.3 | 14.9 | 91.8 | 46.1 | 58.5 | 32.7 | 88.4 | 7.7 | 100 |
| β -sheets | 23.2 | 44.1 | 63.1 | 58.2 | 65.9 | 8.2 | 48.6 | 4.1 | 45.3 | 5.4 | 21.7 | 0 |
| Turns | 33.1 | 0 | 0 | 0 | 19.2 | 0 | 0 | 0 | 22 | 0 | 70.6 | 0 |
| Others | 0 | 0 | 8.3 | 27.5 | 0 | 0 | 5.3 | 37.4 | 0 | 6.2 | 0 | 0 |

*Supplementary Table 3: Secondary structures determination from CD spectra based on K2D2 in peptides grown for 1 week under fibrillation conditions at pH 5.8.*

| | αCoV | βCoV-1 | βCoV-2 | δCoV | γCoV_14 | γCoV_24 |
|---|---|---|---|---|---|---|
| α Helix | 17.4 | 16.4 | 11.3 | 13.0 | 14.6 | 6.3 |
| β Strand | 26.8 | 29.0 | 28.6 | 28.3 | 28.2 | 32.4 |

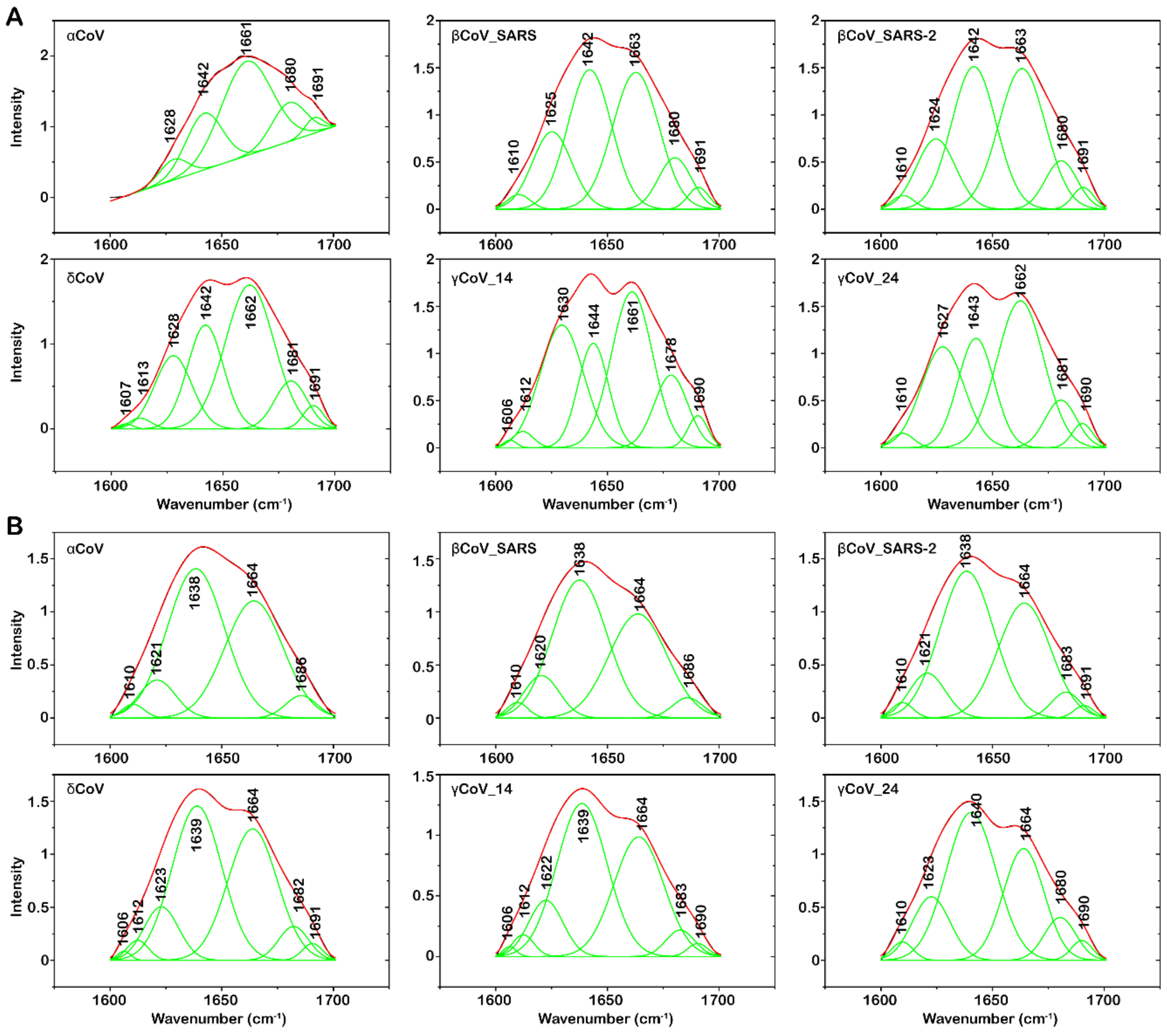


*Supplementary Figure 7. Deconvolution of FT-IR Amide I peak of peptides grown for 1 week under fibrillation conditions at a. pH 5.8 and b. pH 7.5.*

*Supplementary Table 4: Secondary structures identified using FT-IR in peptides grown for 1 week under fibrillation conditions at pH 5.8 and 7.5.*

| | αCoV | | βCoV-1 | | βCoV-2 | | δCoV | | γCoV_14 | | γCoV_24 | |
|---|---|---|---|---|---|---|---|---|---|---|---|---|
| Helix | 60.3 | 38.3 | 40.0 | 37.6 | 40.9 | 35.8 | 49.4 | 36.3 | 41.4 | 34.6 | 43.6 | 31.8 |
| β-sheets | 8.0 | 32.0 | 18.2 | 9.4 | 16.5 | 33.7 | 18.7 | 32.7 | 30.7 | 34.4 | 26.3 | 15.7 |
| Turns | 8.8 | 3.9 | 6.0 | 3.8 | 5.6 | 5.2 | 6.4 | 6.6 | 8.3 | 5.2 | 5.7 | 5.3 |
| Others | 22.9 | 25.8 | 35.8 | 49.6 | 37.0 | 25.3 | 25.5 | 24.4 | 19.6 | 25.8 | 24.4 | 47.2 |
| | pH 5.8 | pH 7.5 | pH 5.8 | pH 7.5 | pH 5.8 | pH 7.5 | pH 5.8 | pH 7.5 | pH 5.8 | pH 7.5 | pH 5.8 | pH 7.5 |

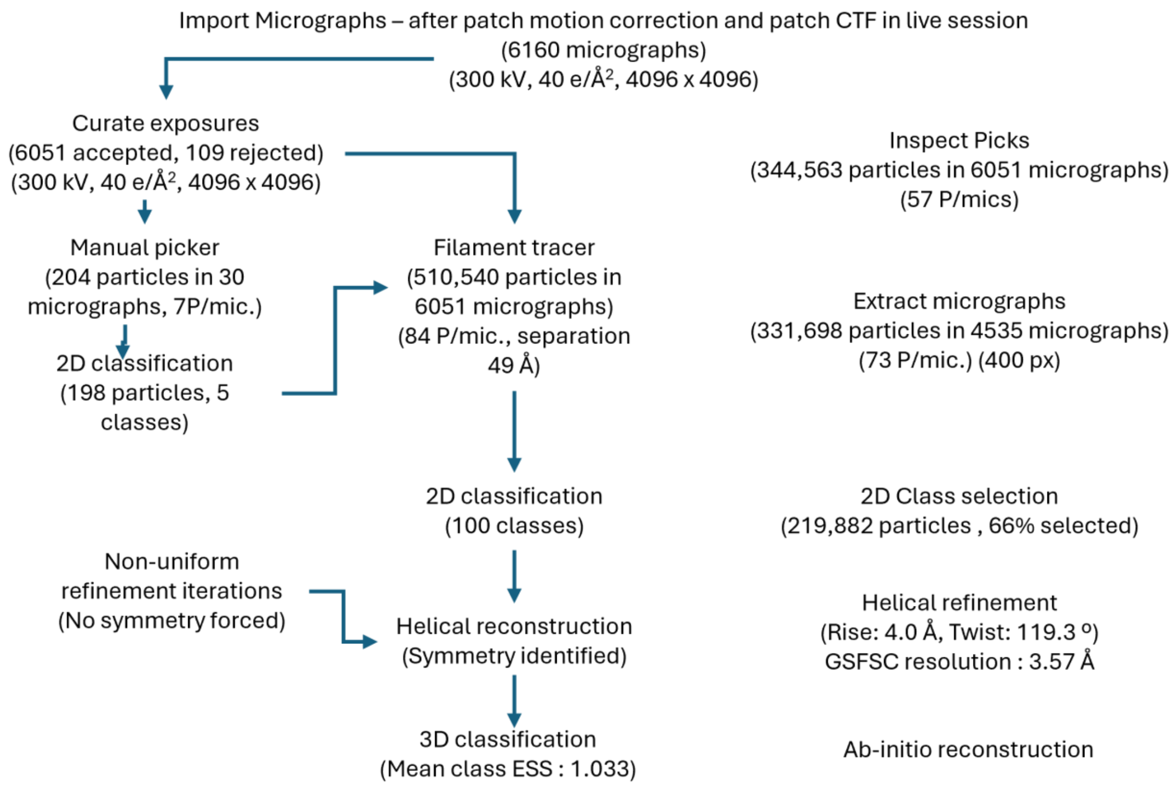


*Supplementary Figure 8. Workflow of cryoEM data processing in CryoSPARC.*

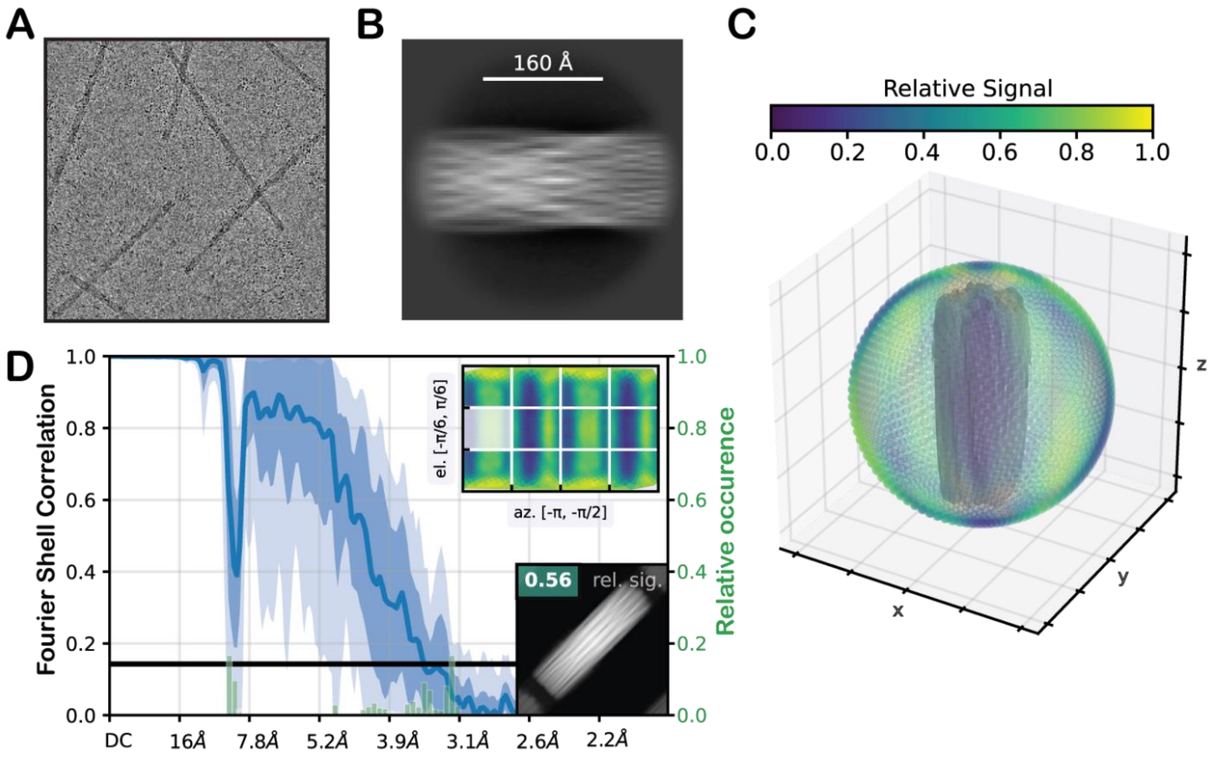


*Supplementary Figure 9. CryoEM model generation and processing workflow in CryoSPARC. (A) Representative cryo-EM micrograph of δCoV fibrils, illustrating predominantly single, well-separated fibrils with minimal overlap. (B) Example of a 2D class average selected for initial model generation.*

*(C) 3D coloured scatter plot showing relative signal as a function of viewing direction, with the corresponding fibril structure embedded at the centre. (D) Fourier Shell Correlation (FSC) curve demonstrating the resolution of the reconstructed 3D model. The top-left inset depicts relative signal distributed across twelve regions of the viewing sphere, defined by azimuth and elevation boundaries. The bottom-right inset shows the model projection along the central viewing direction, along with the mean relative signal for that region.*
